# Ubiquitylome Rewiring by Bacterial E3 Ligases Reveals Multifaceted Host Subversion

**DOI:** 10.64898/2025.12.02.691888

**Authors:** Andrea Bullones-Bolaños, Isela Serrano-Fujarte, Sara Martín-Villanueva, Carla V. Galmozzi, Francine Amaral Piubeli, Jesús de la Cruz, Laura Tomás-Gallardo, Francisco Ramos-Morales, Joaquín Bernal-Bayard

## Abstract

*Salmonella enterica* has evolved an arsenal of effector proteins secreted *via* type III secretion systems (T3SS) to manipulate host cell functions. Among these, the NEL family E3 ubiquitin ligases (SlrP, SspH1, and SspH2) are known to modulate immune signaling, but the breadth of their impact on the host ubiquitylome remains unexplored. In this study, we have performed a global proteomic analysis to identify host proteins ubiquitylated in response to expression of these three effectors in human cells. Using enrichment strategies combined with mass spectrometry under conditions where the proteasome is active or not, we identified 214 putative substrates of ubiquitylation. Gene ontology and KEGG pathway analysis revealed enrichment in pathways related to RNA processing, ribosome biogenesis, cytoskeleton organization, chromatin remodeling, and vesicular trafficking. *In vitro* ubiquitylation assays validated five novel substrates and revealed differential substrate specificity and patterns of ubiquitin chain topology among the effectors. Notably, expression of SspH1 in *Saccharomyces cerevisiae* disrupted polysome profiles in a ligase activity dependent manner, indicating a direct impact of the bacterial effector on translation of eukaryotic cells. Comparison with previously published global interactomes and ubiquitylomes supports a model in which *Salmonella* NEL effectors subvert a broader range of host pathways through targeted ubiquitylation. Our findings uncover new roles for NEL ubiquitin ligases in host manipulation and provide a holistic analysis of their effects on the ubiquitylome of the host cell, constituting a valuable resource for the study of bacterial pathogenesis and infection biology.

## INTRODUCTION

Nontyphoidal *Salmonella enterica* in one of the most important intracellular pathogens, responsible for an estimated 93.8 million of cases of gastroenteritis and over 150,000 deaths annually ^1^. As other Gram-negative bacteria, *Salmonella enterica*, have evolved sophisticated type III secretion systems (T3SS) to inject effector proteins directly into eukaryotic host cells ^2^. These effectors manipulate host signaling pathways and cellular processes, enabling bacterial survival and proliferation ^3^. The ubiquitylation system is a crucial post-translational modification in eukaryotic cells that regulates a wide range of cellular processes, including protein degradation, signal transduction, membrane protein trafficking, and immune responses ^4^. In the context of infections, the ubiquitylation system plays a fundamental role in the host immune response, regulating the activation of inflammation, selective autophagy, and programmed cell death to eliminate invading pathogens ^5^. However, many intracellular pathogens, including *Salmonella*, have the capacity to manipulate the host ubiquitylation system to their advantage ^5,6^. For instance, some bacteria express effector proteins that act as E3 ubiquitin ligases ^7,8^ or deubiquitinases (DUBs) ^9,10^, mimicking or interfering with the host ubiquitylation machinery. The NEL family of E3 ligases is a unique group of bacterial enzymes that do not share sequence homology with eukaryotic E3 ligases ^11^.

Several NEL bacterial E3 ligases have been identified in *Salmonella enterica*, including SlrP, SspH1, and SspH2. These effectors are characterized by an N-terminal domain involved in T3SS-dependent translocation, a central leucine-rich repeat (LRR)-containing LPX domain thought to mediate protein-protein interactions, and a C-terminal NEL domain responsible for E3 ligase activity. A limited number of host substrates have been identified for these effectors to date: SlrP modifies the essential redox protein Thioredoxin ^12^, and the spliceosomal component SnrpD2 ^13^; SspH1 targets PKN1, a serine/threonine kinase involved in NF-κB and androgen receptors signaling ^11,14^; and SspH2 ubiquitinates Nod1, an intracellular pattern recognition receptor that detects the components of the bacterial cell wall and activates the innate immune response ^15^. More recently, SspH2 has been shown to target the LIM-domain scaffold protein LMO4 for proteasomal degradation, suggesting interference with IL-6-mediated STAT3 activation ^16^. Together, these findings highlight the immune-modulatory potential of NEL ligases at multiple signaling levels. However, it is believed that there are many more substrates for *Salmonella* NEL effectors ^17^. Defining these targets is crucial to elucidate how *Salmonella* rewires host signaling networks in order to promote intracellular survival and pathogenesis.

In this study, we have used a combination of ubiquitin enrichment techniques coupled to a semi-quantitative proteomic analysis to investigate the ubiquitylome of HEK293T cells expressing SlrP, SspH1, and SspH2. This approach has allowed us to identify up to 214 novel potential host targets for *Salmonella* NEL effector-mediated ubiquitylation. Our findings suggest that SlrP, SspH1, and SspH2 target a variety of host processes, including RNA metabolism, protein translation and modification, and cytoskeleton dynamics. Furthermore, we have confirmed the *in vitro* ubiquitylation of five of these host proteins by the *Salmonella* NEL effectors and have revealed the specificity and the potential functional redundancy of SlrP, SspH1, and SspH2 towards these proteins. Finally, we have focused on SspH1, whose expression in yeast, not only produces toxicity to the cells, but also causes an alteration in ribosome function, specifically dependent on its ubiquitin ligase activity, suggesting that SspH1 impacts protein synthesis.

Understanding how pathogens manipulate host processes at the molecular level is essential for the development of novel therapeutic strategies to combat bacterial infections. Together, our findings revealed a broad and previously underappreciated role for *Salmonella* NEL ligases in the manipulation of host gene expression, from transcription to translation. By uncovering novel substrates and mechanistic insights, this study provides a valuable resource for the field and lays the groundwork for future efforts to dissect effector-substrate interactions in the context of infections.

## RESULTS

### Heterologous expression of Salmonella enterica NEL effectors modifies the host ubiquitylome

To analyze the impact of *Salmonella* NEL effectors on the host ubiquitylation system, we performed a semi-quantitative proteomic analysis of ubiquitylated host proteins in presence of SlrP, SspH1 and SspH2. Our approach combined ubiquitin-protein enrichment coupled to LC-MS/MS, enabling us to identify the pool of host-cell protein targets that are ubiquitylated by the three NEL ubiquitin-E3 ligases encoded in the genome of *Salmonella enterica* serovar Typhimurium (**Figure 1A**). Briefly, HEK293T cells were transiently transfected either with pcDNA3 derivative vectors simultaneously expressing SlrP, SspH1 and SspH2 or with the pcDNA3 empty vector. Considering that proteasomal degradation could represent one of the primary fates of the ubiquitin-tagged proteins ^18^, we additionally performed the experiment in the presence of the proteasome inhibitor MG132, in order to prevent potential loss of putative effector substrates due to degradation. To ensure that correct heterologous expression of each effector took place in the transfected cells, we took advantage of a FLAG epitope introduced at the C-terminus of each effector (**Figure S1 C-D**). Cells were harvested 24 h after transfection, lysed and total protein was quantified in the cell extracts. Upon quantification (**Table S3**), the UBIQAPTURE-Q kit (Enzo life Sciences) was used to perform ubiquitylated protein enrichment of each sample **(Figure S1 A-B**). This system includes a high affinity matrix for ubiquitin monomers and polymers. The ubiquitylated proteins were trypsin-digested and identified by LC-MS/MS. The mass spectrometry dataset was analysed with the Thermo Proteome Discoverer v2.2.0.388 (PD) (Thermo Scientific Inc.) Software tool and the Uniprot-human-Reference database (UP000005640) was used for protein identification. Search parameters included fixed modifications such as carbadomethylations, as well as variable modifications such as GlyGly (K), LRGG, oxidation (M), RGG (K), and N-terminal acetylation. A minimum peptide length of seven amino acids was required and the false discovery rate (FDR) at the protein level was set at 0.02. Putative substrates of the three NEL ubiquitin E3 ligases were identified using stringent selection criteria (log_2_FoldChange >2.5 and -log_10_pValue >1.3). The resulting candidate proteins for each experimental condition are summarized in **Table S1**. Although all proteins meeting these thresholds were considered potential substrates, single-hit identifications need additional validation. To establish a high-confidence substrate pool, we performed comparative proteomics between control samples (982 total proteins) and effector-expressing samples (980 total proteins) under both standard (- MG132) and proteasome-inhibited (+ MG132) conditions. Biological replicates exhibit high reproducibility, with Pearson’s correlation coefficient of up to r = 0.96 values (**Figure 1B, 1E, and Figure S2**) and consistent abundance distributions across independent experiments (**Figure S2**). Differential enrichment analysis revealed significant accumulation of ubiquitylated proteins in the presence of *Salmonella enterica* E3 ligases compared to control conditions (**Figures 1C and 1F**), suggesting that these bacterial effectors actively modify host protein ubiquitylation patterns. We were able to generate a list of potential host cell substrates for the *Salmonella* NEL effectors identified by mass-spectrometry from both conditions, MG132 -/+, then we built two separate ubiquitylated-substrate networks, containing 110 and 104 proteins, respectively (**Figure 1D, 1G, and Table S1**). STRING-based interaction network analysis showed consistent clustering of identified targets into functional modules. Several biological processes were significantly enriched in both experimental conditions, most notably RNA processing and translation, suggesting conserved cellular hubs preferentially targeted by *Salmonella* NEL ligases. However, condition-specific enrichment patterns were also observed. In the absence of MG132, we identified clustering of proteins involved in cytoskeletal polymerization, mitochondrial gene expression, and programmed cell death. In contrast, under MG132 treatment, we observed a marked enrichment in proteins associated with post-translational protein modification, suggesting a global manipulation of the ubiquitylation system by the effectors. The exclusive appearance of these proteins when the proteasome is inhibited suggests that they might be rapidly degraded after ubiquitylation, which would explain their absence in the condition without the proteasome inhibitor. These findings suggest coordinated targeting of core host processes, supporting the existence of new, previously undescribed, roles for *Salmonella* NEL ligases beyond innate immunity.

**Figure 1.**
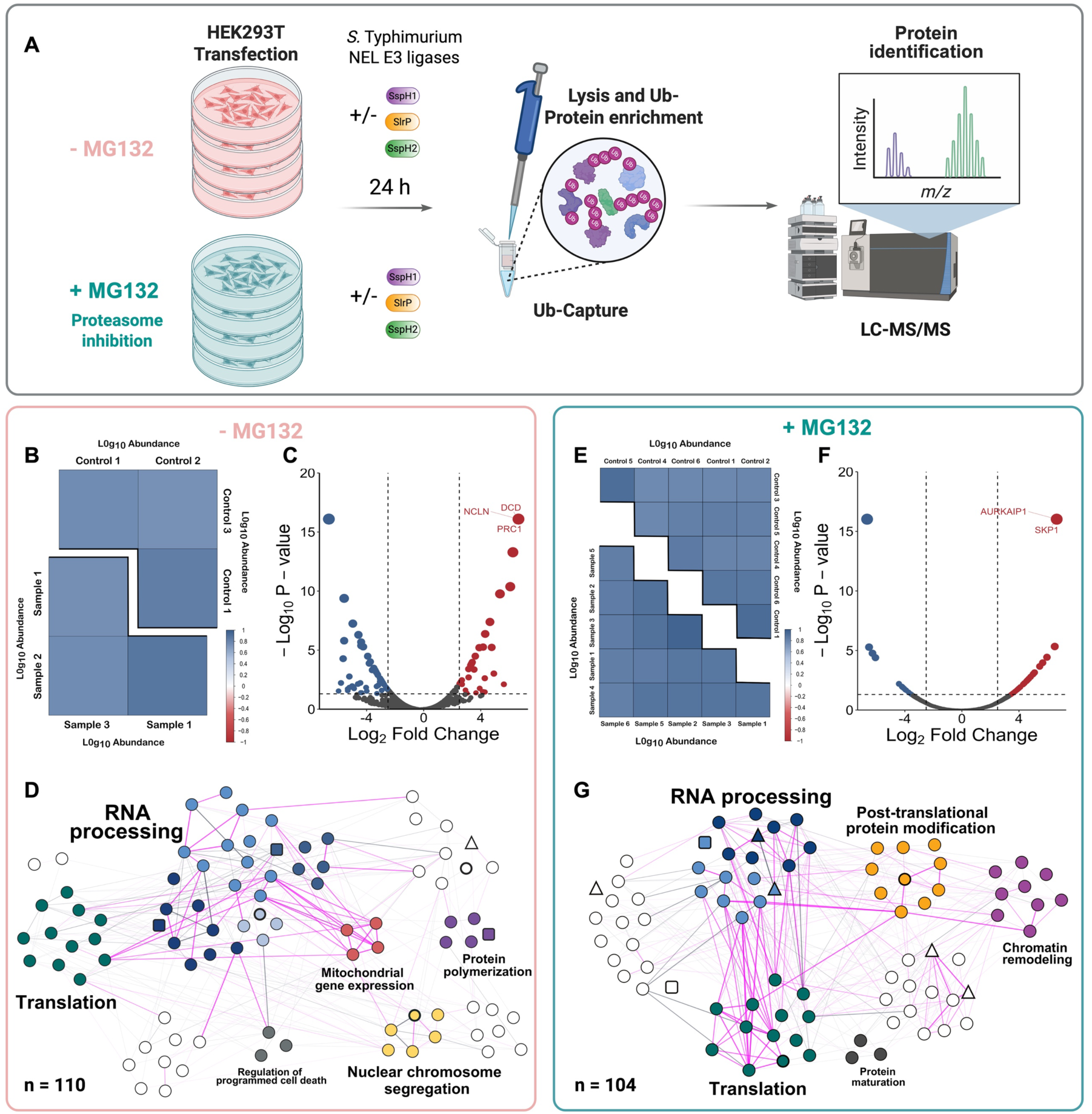
Pipeline for enrichment and analysis of ubiquitylated host substrates by *Salmonella* NEL effectors. (A) Empty pcDNA3 or derived plasmids expressing NEL effectors were used to transfect HEK293T cells. Samples were treated or not with the proteasome inhibitor MG132 for 6 h. Total cell extracts were subjected to mono- and poly-ubiquitylated protein isolation using a high binding affinity matrix. Captured proteins were trypsin-lysed and run in an Orbitrap Q-Exactive Plus mass spectrometer. (B and E) Pearson correlation analysis of non-scaled protein abundances between biological replicates, the change of intensity in the color scale shows the increase or decrease in r values. The scatter plots can be found in Figure S2. (C and F) Volcano plot visualization of differential protein enrichment comparing control and NEL-effector expressing samples. Vertical dashed lines indicate significance thresholds (log_2_ Fold Change > 2.5 or < −2.5), horizontal dashed line represents statistical significance threshold (-log_10_pValue >1.3). Significantly enriched proteins (red) represent putative NEL ligase substrates, while significantly depleted proteins are shown in blue. Gray points indicate proteins that did not meet significance criteria. The size of the dots represents an increasing number of proteins with identical criteria (see Table S1). (D and G) STRING-based protein interaction network of NEL-targeted host proteins identified in this study. Nodes represent individual proteins and are shaped as follows: circles indicate novel substrates, squares represent protein previously identified in other studies, and triangles represent members of protein families previously identified. Nodes are colored according to their assigned biological processes (corresponding to classification in Figure 2A and 2D). Edges represent protein-protein interactions, with pink edges highlighting experimentally validated interactions according to the STRING database. The validated proteins are highlighted with bold outlines. n = number of proteins identified in each experimental condition.

### Salmonella NEL effectors impact diverse host processes

The host ubiquitylation machinery plays a crucial role regulating a wide range of cellular processes, from the classical role of proteasomal degradation to DNA repair, endocytic trafficking, and cell cycle progression ^19^, as well as modulation of the immune system ^20^ and pathogen clearance ^21^. On the other hand, *S. enterica* manipulates multiple cellular functions and modulates the host defense systems to its advantage during infection, employing a diverse arsenal of virulence factors encoded within its genome ^3^. To gain insight into host cellular processes specifically enriched in the presence of the *Salmonella* NEL ubiquitin ligases, both under proteasome inhibition and non-inhibition conditions, we performed a Gene Ontology (GO) enrichment analysis using the pool of potential substrates identified by mass spectrometry. RNA processing and translation appear to be the most enriched processes in both experimental conditions (**Figure 2A and 2D**). Among subcategories of RNA processing, ribonucleoprotein complex and ribosome biogenesis, RNA catabolic process and RNA splicing are overrepresented, suggesting that *Salmonella enterica* may manipulate the host gene expression program through the ubiquitylation machinery. GO analysis also categorized the ubiquitylated targets by cellular component, highlighting cell structures like ribosome, nucleus, ubiquitin machinery, focal adhesion or cytoskeleton as the most significant. Interestingly, SspH1, which has been reported to enter the host nucleus ^22^, has been classified as a nucleomodulin ^23^, and represses NF-κB-dependent gene expression, downregulating the host inflammatory response ^14^. Moreover, SlrP targets and ubiquitylates SnrpD2, one of the essential components of the splicing machinery ^13^. Other studies have identified these or functionally related ligands, like RBM proteins, pointing to splicing as a target process of SlrP ^24^. Ubiquitylation affects ribosome biogenesis by influencing the expression, assembly and degradation of ribosomal proteins, as well as the function of ribosomal biogenesis factors and the elimination of mature ribosomes ^25^. In this line, the “mitochondrial gene expression” GO category includes 4 proteins of the MRP (mitochondrial ribosomal protein) family that constitute structural elements of the mitochondrial ribosome and contribute to translation. Interestingly, the GO group “nuclear chromosome segregation” includes host proteins like the SMC2/4 subunits of the condensin complex, the chromatin remodeler CHD3, or UHRF, which is a E3 ubiquitin ligase that acts as epigenetic regulator by bridging DNA methylation and chromatin modification. Ubiquitylation of these proteins could modulate host gene expression. PRC1, experimentally validated (**Figure 3**), ECT2 and CUL1 are involved in cell cycle progression. Interestingly, other over-represented cellular processes that are in line with what other authors have identified in the context of *Salmonella* pathogenesis are protein polymerization ^26^, and regulation of programmed cell death ^12^. *Salmonella* employs a multifaceted strategy to manipulate actin polymerization, using diverse effectors, that act at different stages of infection to promote invasion, intracellular survival, and modulation of host responses ^26^. SspH2 interacts with key proteins that regulate actin dynamics and can inhibit actin polymerization *in vitro* ^27^. In addition, several actin cytoskeleton-related proteins, such as ACTR8, ARPC5, MYH9, SUGT1, and DYNC1H1 have been identified as binding partners ^24^, suggesting that SspH2 could manipulate the host actin network through direct interactions, while none of these ligands has been previously shown to be substrates of its E3 ligase activity. Our ubiquitylome analysis recapitulates several previously identified targets or members from the same family like DYNLL2 or MYO1C, as well as ACTR2/3, from the ARP2/3 complex, which plays a key role in the actin organization required for bacterial internalization ^28^. Related to that, the mass spectrometry data set also includes putative substrates of NEL effectors that are involved in intracellular vesicular trafficking, such as, Rab14, the Ras-related protein R-Ras2, and phosphatidylinositol 4-kinase (PI4K). These proteins play critical roles in endocytic recycling, membrane dynamics, and phosphoinositide signaling, suggesting that *Salmonella* NEL ligases could modulate trafficking pathways to support bacterial survival and replication.

**Figure 2.**
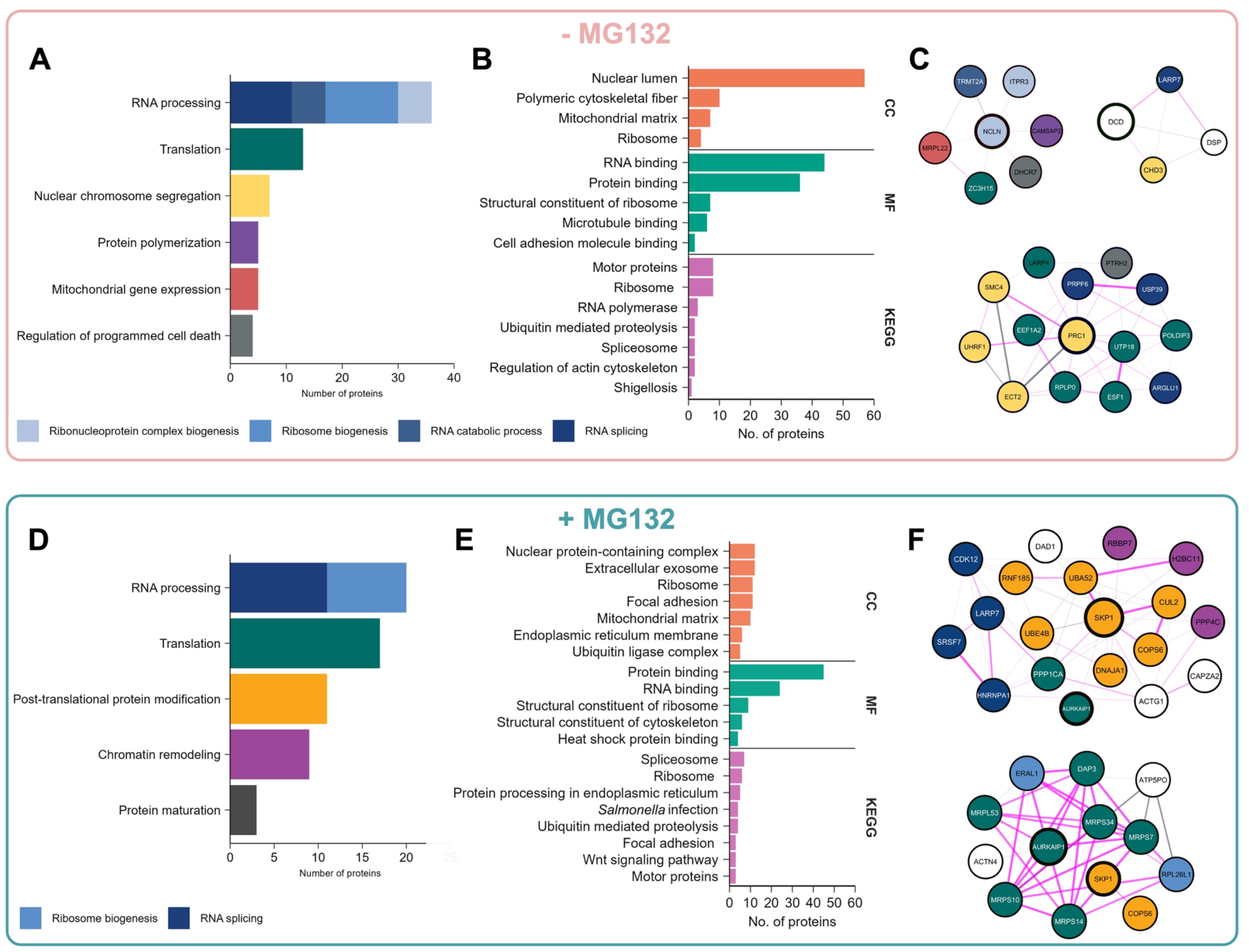
Enrichment analysis of host proteins targeted by *Salmonella* NEL effectors. (A and D) Enrichment analysis of biological processes associated with NEL-targeted proteins. (B and E) Multi-category enrichment analysis showing the distribution of NEL-targeted proteins across cellular components (orange upper bars), molecular functions (green middle bars), and KEGG pathways (pink bottom bars). (C and F) STRING-based interaction networks of experimentally validated NEL effector substrates with the other substrates identified in the study. Nodes represent individual proteins and edges indicate protein interactions (experimentally validated interactions according to the STRING database, are shown in pink). Colors denote their biological process according to graphs A and D, respectively. The validated proteins are highlighted with bold outlines.

**Figure 3.**
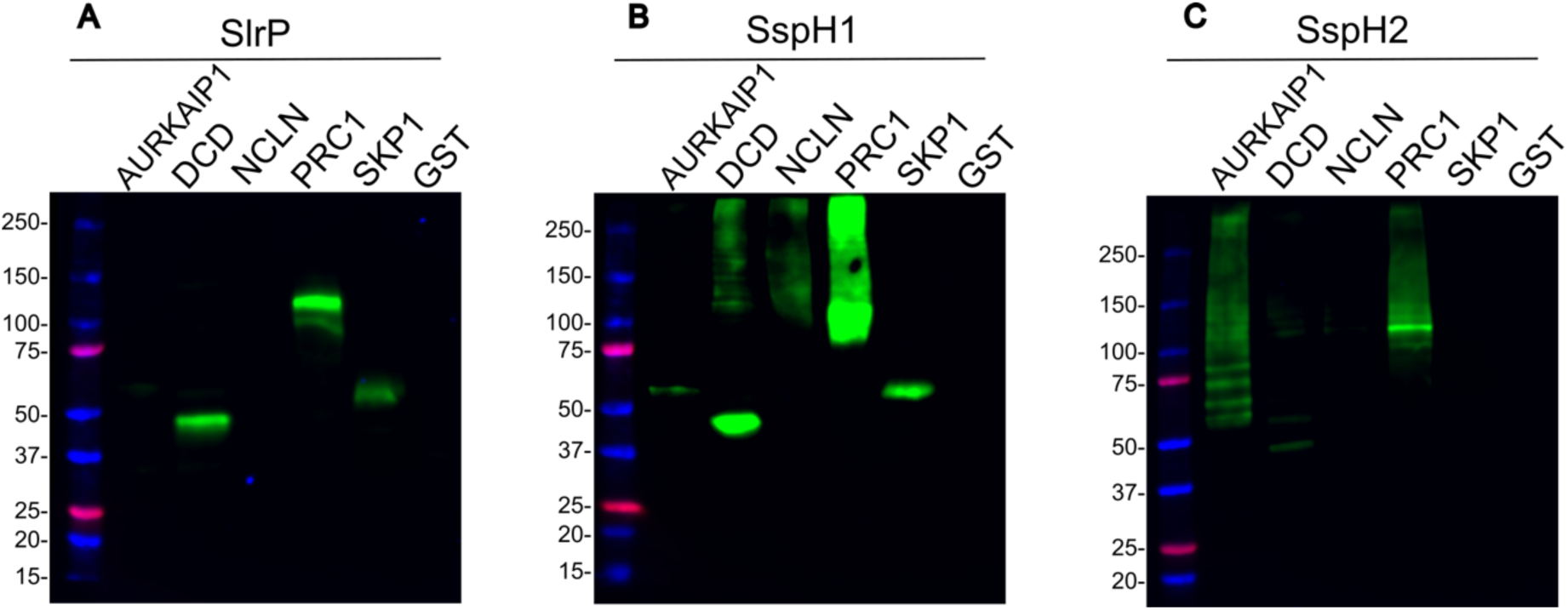
Validation of newly identified *Salmonella* NEL substrates by *in vitro* ubiquitylation reactions. Ubiquitylation reactions were carried out with GST fusion proteins bound to glutathione-agarose beads, in the presence of E1, E2, HA-ubiquitin and the effectors SlrP (A), SspH1 (B) and SspH2 (C). Beads were washed before immunoblot analysis with anti-HA monoclonal antibodies. The results shown are representative of three independent experiments.

Our results are also consistent with the KEGG pathways analysis, that revealed distinct processes targeted by the NEL family effectors (**Figure 2B and 2E**). Notably, proteins in pathways related to *Salmonella* and *Shigella* infection, and actin polymerization were found, suggesting that these effectors could be involved in the manipulation of the host systems at different stages of infection. In both conditions, pathways related to RNA processing (splicing, ribosome) are enriched, indicating that NEL ligases strategically target the host translation machinery, potentially to redirect cellular resources toward bacterial replication or to inhibit synthesis of relevant host defense factors. The KEGG pathways also include proteins that are involved in ubiquitylation processes consistent with the ubiquitin ligase activity of these effectors. In summary, this targeted modification represents a sophisticated virulence strategy, as these pathways represent a central hub, connecting the various cellular functions mentioned earlier and relevant molecular processes, as shown in **Figure 2C and 2F**. Collectively, our proteomic analysis shows that multiple host cellular functions are hijacked during *Salmonella* infection via NEL dependent ubiquitylation, and the *Salmonella*-induced modification of the ubiquitylation landscape described here represents a valuable resource for the scientific community to explore new mechanism underlying bacterial pathogenesis.

### Ubiquitylation *in vitro* validates the identification of new host cell substrates for Salmonella NEL effectors

One limitation of the approach used in this study is that we are unable to directly assign specific substrates to each of the *Salmonella* NEL ligases. However, with the compiled list of potential candidates (**Table S1**), it is possible to evaluate the potential redundancy and specificity of the effectors towards each substrate. This can be achieved by *in vitro* ubiquitylation assays, which simultaneously serve to validate the efficiency of the Ubi-capture coupled to mass spectrometry strategy.

For experimental validation, we selected 5 of the most significantly enriched ubiquitylated proteins from our data set from both experimental conditions, with and without MG132 (**Figure 1C and 1F**). Among the candidates from samples not treated with MG132, DCD (Dermcidin), NCLN (Nicastrin-Like Protein) and PRC1 (Protein regulator of cytokinesis 1) were selected, while AURKAIP1 (Aurora Kinase A Interacting Protein 1) and SKP1 (S-phase kinase-associated protein 1) were picked from MG132-treated samples. The chosen proteins are involved in several biological processes enriched according to our interaction network analysis (**Figure 1D, 1G**), including mitochondrial protein translation (AURKAIP1), progression of cell cycle (PRC1), and ubiquitylation (SKP1), or could have an interesting function in the context of infection, like NCLN, involved in ER protein transport, or DCD, which display antimicrobial activity.

Genes encoding the selected candidates were subcloned in the pGEX derivative vector to produce GST-tagged purified versions of each protein and used for *in vitro* ubiquitylation assays. First, SlrP (**Figure 3A**) was shown to ubiquitylate DCD, PRC1 and SKP1. This effector only adds one ubiquitin to these substrates as we observed a single band roughly matching the molecular weight of the protein (GST-DCD ∼37 kDa, GST-PRC1 ∼98 kDa, GST-SKP1 ∼44 kDa) plus ∼10 kDa corresponding to the HA-tagged ubiquitin. On the other hand, SspH1 (**Figure 3B**) was also able to ubiquitylate these three substrates (DCD, PRC1 and SKP1), but unlike SlrP, it produced polyubiquitylation of DCD and PRC1. Additionally, SspH1 is the only one of the three effectors capable of polyubiquitylating NCLN (GST-NCLN ∼87 kDa). SspH2 (**Figure 3C**) produced ubiquitin chains specifically on AURKAIP1 (GST-AURKAIP1 ∼49 kDa), although it also ubiquitylated DCD and PRC1, but not SKP1 or NCLN. These *in vitro* ubiquitylation analysis validate the selected candidates as substrates of the ubiquitin ligase activity mediated by these bacterial effectors. Importantly, it also allowed us to identify new specific substrates for SspH1 (NCLN) and SspH2 (AURKAIP1). Additionally, these assays revealed the existence of certain degree of specificity and redundancy among *Salmonella* NEL effectors, both in substrate selection and in the type of modification they perform, as substrates such as DCD and PRC1 are common to all three E3 ligases, although with different forms of modification (SlrP: monoubiquitylation, SspH1 and SspH2: polyubiquitylation). Though we observed redundancy among some of the newly identified substrates of the *Salmonella* E3 ligases, the fate of the ubiquitylated proteins by each effector could be different since, as we demonstrated in a previous work, while SspH1 and SspH2 preferentially catalyze K48-linked ubiquitin chains (related to protein degradation by proteasome), SlrP performs other types of ubiquitin linkage than K48 ^29^.

### First-neighbor interaction networks showed diverse functional connectivity among NEL substrates

To understand the possible roles of the validated novel NEL-substrates, we built the first-neighbor interaction subnetworks and functionally annotated the proteins identified (**Figure 2C and 2F**). In general, the highest enriched biological processes across the networks of these five new NEL-substrates were: translation, RNA splicing, and post-translational modification. It also showed extensive interconnectivity among distinct biological processes for each selected candidate.

Among the five validated targets, NCLN (**Figure 2C**), classified within “ribonucleoprotein complex biogenesis” biological process, showed connections with six immediate interactors spanning diverse functional categories, including protein polymerization, cell death regulation, translation, RNA splicing, and mitochondrial gene expression. This connectivity pattern led us to hypothesize that NCLN could be targeted due to its potential role as a linker between three relevant processes during bacterial infection: RNA processing machinery, cytoskeletal dynamics, and cellular stress responses.

DCD, although lacking a defined functional annotation (**Figure 2C**), showed interactions with proteins involved in nuclear chromosome segregation and RNA splicing, like Larp7, which has been described to be a target for intracellular pathogens ^30^. Notably, one of its interactors, DSP, although in our analysis lacks biological process annotation, is implicated in filament organization ^31^, and this is a pathway exploited during *Salmonella* infection of epithelial cells ^32^.

PRC1 shows extensive connectivity, with 14 first-neighbor interactions predominantly comprising translation-associated factors (6 proteins) and RNA splicing components (3 proteins); the additional connection to programmed cell death regulation (1 protein) revealed an unexpected link between chromosomal dynamics, translational control, and possibly stress responses (**Figure 2C**).

SKP1, involved in post-translational protein modification (**Figure 2F**), showed the largest network, with 18 interactors spanning diverse functional categories: its own process (6 proteins), chromatin remodeling (3 proteins), translation (2 proteins), and RNA splicing (4 proteins). The significant representation of chromatin remodelers suggests coordination between ubiquitylation machinery and epigenetic regulation, potentially enabling broad transcriptional reprogramming through targeted modifications. The last validated NEL target, AURKAP1, was interconnected with 12 more proteins, most of them involved in translation (like AURKAP1 itself), post-translational modification components (2 proteins), and ribosome biogenesis factors (2 proteins). Remarkably, we identified reciprocal interactions between SKP1 and AURKAP1, both found in each other’s subnetwork (**Figure 2F**), establishing a link between translation and post-translational modification systems that may represent a strategic vulnerability exploited by these *Salmonella* E3 ubiquitin ligases.

### SspH1 impacts translation in Saccharomyces cerevisiae

According to our proteomic results, the expression of SlrP, SspH1, and/or SspH2 might target essential host cellular functions like ribosome biogenesis and protein translation (**Figure 1D, 1G** and **Figure 2A, 2D**). Importantly, ubiquitylation plays a significant role in ribosome biogenesis, by regulating the stability of ribosomal proteins ^25^. Thus, many ribosomal proteins are subjected to rapid degradation via ubiquitylation by the proteasome when they do not readily assemble into functional ribosomal subunits ^33,34^. In addition, other studies have demonstrated that ubiquitylation of well-assembled ribosomal proteins also plays distinct regulatory roles in translation control ^25,35,36^. Thus, to address the *in vivo* impact of *Salmonella* ubiquitin ligases on ribosome assembly and translation, we used the gold-standard model organism for the study of these two processes in eukaryotes ^37–39^: the yeast *Saccharomyces cerevisiae.* We first assessed growth in yeast expressing each of the three bacterial effectors (**Figure 4A**). To this end, we generated pAS24-based plasmid constructs encoding each effector under the control of a galactose-inducible promoter. To verify the expression of the bacterial genes in yeast, C-terminal HA-tagged versions of the effectors were used (**Table S2**, **Figure 4B**). When transformants were grown on repressive glucose-containing solid media, their growth was indistinguishable from that of the control transformed with the empty pAS24 plasmid. In contrast, on galactose-containing media, yeast growth was strongly inhibited by SspH1 expression, while expression of SlrP and SspH2 had no observable effect. Interestingly, the catalytically inactive mutant SspH1^C492A^ failed to inhibit yeast cell growth (**Figure 4A**), indicating that this effect on growth is mediated by ubiquitylation. These results, consistent with previous work ^40^, indicate that SspH1 might target and ubiquitylate a critical yeast substrate required for cell growth. To address whether this growth defect is related to ribosome biogenesis and/or translation, yeast cells were transformed with the pAS24 empty plasmid or pAS24 constructs harboring the wild-type SlrP, SspH2, and SspH1 alleles or the mutant SspH1^C492A^ variant, and initially cultivated in the presence of raffinose as a carbon source. Then, galactose was added to fully induce the expression of these different proteins for 16 h (**Figure 4B**) and polysome profiles were analyzed. As shown in **Figure 4C**, cells transformed with the empty pAS24 plasmid exhibited the characteristic wild-type polysome profiles, displaying normal levels of free 40S and 60S ribosomal subunits, as well as intact polysomes. Similar profiles were observed for the yeast cells expressing SlrP and SspH2 (**Figure 4D and 4E**). In clear contrast, expression of SspH1 revealed a noticeable reduction in the overall polysome content, as revealed by the profiles shown in **Figure 4 F.** This profile, commonly observed in many translation initiation mutants ^41^, is indicative of a translation initiation defect. None of the profiles showed the characteristic features of a defect on the biogenesis of 60S or 40S ribosomal subunits, such as half-mer polysomes or an imbalance in free 40S and 60S subunits ^42^. Expression of the catalytically dead SspH1^C492A^ allele generates a normal polysome profile, as it was entirely comparable to those of cells harboring the empty vector **(Figure 4G**). Altogether, these results indicate that the ubiquitylation activity of SspH1 significantly impairs translation initiation in yeast.

**Figure 4.**
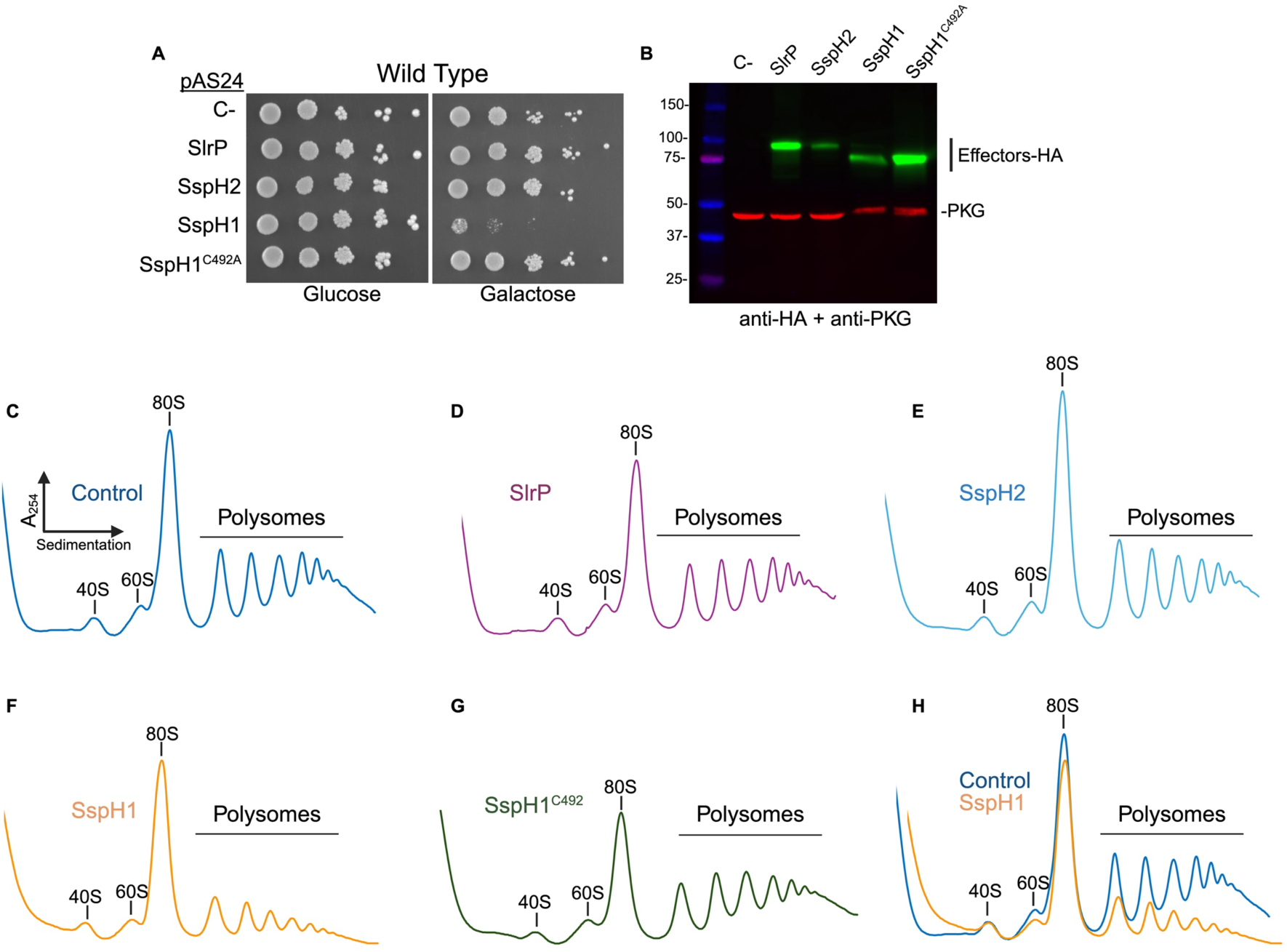
SspH1 impairs translation initiation in *S. cerevisiae*. (A) SspH1 expression affects yeast growth. Wild type yeast cells carrying pAS24 derivative plasmids encoding galactose-inducible SlrP, SspH1, SspH2 and SspH1^C492A^ were spotted in 10-fold serial dilution steps onto minimal medium plates without leucine and glucose or galactose as a carbon source. Plates were incubated at 30 °C for 3 days. (B) Heterologous expression of NEL effectors in yeast. Strains from panel A were grown in liquid minimal medium in the presence of raffinose. Then, galactose was added to induce the different proteins for 16 h. Total protein extracts were performed and loaded in SDS-PAGE. Then, the presence of the effectors was detected by immunoblotting with an anti-HA antibody. Anti-PGK antibody was used as loading control. (C-G) SspH1 expression alters polysome profiles. Wild type yeast cells carrying pAS24 derivatives were grown in the presence of raffinose. Then, galactose was added to fully induce the different proteins for 16 h. Whole cell extracts were prepared and polysome profiles were performed as indicated in Materials and Methods. In all cases, 10 A_260_ units of each extract were resolved in 10-50% sucrose gradients. The A_254_ was continuously measured. Sedimentation is from left to right. The peaks of free 40S and 60S r-subunits, vacant 80S ribosomes/monosomes and polysomes are indicated. (H) Overlap of polysome profiles of control strain and SspH1 expressing yeast.

### SspH1 Impairs Translation Initiation via the eIF4E-Binding Protein Eap1

To identify the mechanism responsible for the impairment of translation initiation caused by SspH1, we explored the possible involvement of established cellular regulators known to mediate general inhibition of translation initiation. There are two major mechanisms that regulate inhibition of translation initiation in yeast. One of them involves the phosphorylation of the translation initiation factor eIF2α by the non-essential protein kinase Gcn2 ^43,44^. Gcn2 is activated in response to different cellular stresses, most notably amino acid starvation. It has been well reported that deletion of *GCN2* bypasses the stress-induced translational inhibition exerted by eIF2α, as it can no longer be phosphorylated in the absence of this kinase ^44^. To test whether the observed translation inhibition by expression of SspH1 relies on activation of Gcn2, we first analyzed the growth of a *gcn2* null strain expressing SspH1. As shown in Figure 5A, the growth defect caused upon expression of SspH1 was not abrogated by the *GCN2* deletion, making unlikely a role of Gcn2 in the host response to SspH1 expression. Consistently, polysome profiles of *gcn2* null cells were similar to those of wild-type cells upon SspH1 induction (**Figure 5B**).

**Figure 5.**
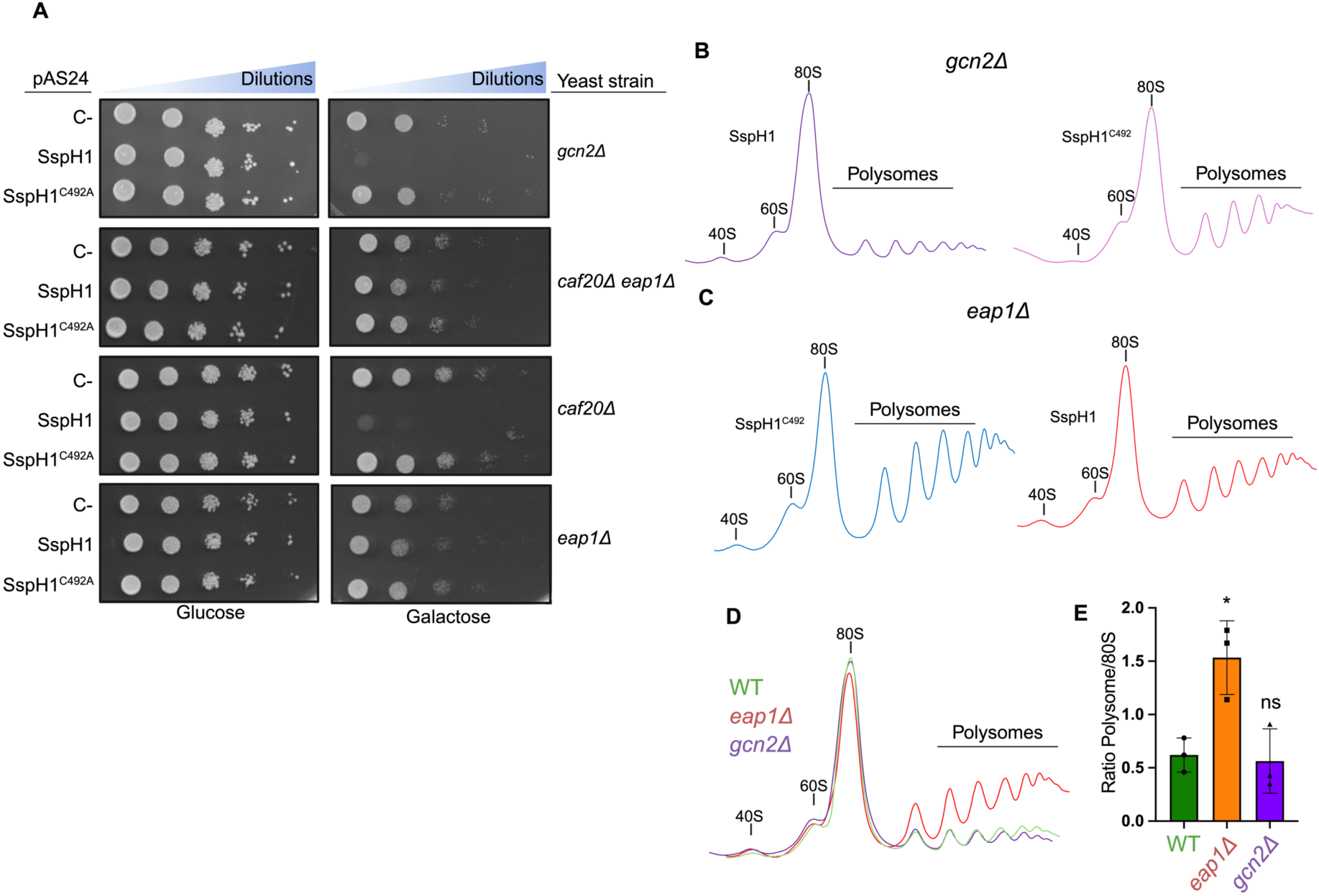
The yeast 4E-binding protein Eap1 mediates the translation inhibition induced by the SspH1 effector. (A) An *eap1* null strain is insensitive to SspH1 expression. The *gcn2*, *cap20 eap1*, *caf20* and *eap1* null mutants were transformed with empty pAS24 or pAS24 derivative plasmids encoding galactose-inducible SspH1 and SspH1^C492A^. Selected transformants were spotted in 10-fold serial dilution steps onto minimal medium plates without leucine and glucose or galactose as a carbon source. Plates were incubated at 30 °C for 3 days. Deletion of EAP1 suppresses the translation inhibition response leads by SspH1. Isogenic wild-type and *eap1* null yeast cells carrying pAS24 derivatives SspH1, and SspH1^C492A^ (B, C) were grown in the presence of raffinose. Then, galactose was added to fully induce the different proteins for 16 h. Whole cell extracts were prepared and polysome profiles were performed as indicated in Materials and Methods. In all cases, 10 A_260_ units of each extract were resolved in 10-50% sucrose gradients. The A_254_ was continuously measured. Sedimentation is from left to right. The peaks of free 40S and 60S r-subunits, vacant 80S ribosomes/monosomes and polysomes are indicated. (D) Overlay of polysome profiles from wild-type, *eap1Δ*, and *gcn2Δ* cells expressing SspH1. The experiments were repeated at least three times, and a representative result is shown. (E) The area under the curve of the yeast polysomes of the different strains expressing SspH1 was quantified, and the ratio between the area of the polysomes and the monosomes was calculated. Bars represent the mean of 3 independent experiments, and individual data points are shown. Statistical analysis corresponds to two tailed unpaired *t-test* with Welch correction. \**p<0.05*. Error bars represent SD.

A second major mechanism regulating the inhibition of translation initiation in eukaryotes involves the hypophosphorylation of the so-called eIF4E-binding proteins (4E-BPs). In their hypophosphorylated state, 4E-BPs bind to eIF4E, which leads to the inhibition of cap-dependent translation initiation ^45^. In yeast, there are two 4E-BPs, namely Caf20 and Eap1, which are both non-essential, and partially redundant ^46,47^. To determine whether the effect of SspH1 on translation initiation relies on the function of Caf20 and/or Eap1, we analyzed the effect of expressing SspH1 in a double *caf20 eap1* null mutant as well as in the corresponding single *caf20* and *eap1* null mutants. As shown in **Figure 5A**, deletion of *EAP1* but not that of *CAP20* clearly alleviates the growth impairment caused by expression of SspH1, thus suggesting that Eap1 could be involved in mediating the translation inhibition response triggered by expression of SspH1. To confirm this hypothesis, we obtained polysome profiles from *eap1* null cells expressing SspH1 and compared them with those obtained from wild-type cells in the same conditions. As shown in **Figure 5C**, deletion of EAP1 yield almost wild-type profiles, as evidenced by the increase in the peaks of polysomes. Overlay of the polysome profiles shown in **Figure 5D** revealed a clear recovery of the polysome peaks in the *eap1* mutant, while both wild-type and *gcn2* strains exhibited a marked collapse of polysomes in the presence of SspH1, indicative of impaired translation initiation. These findings were substantiated by quantitative analysis of the ratio of the Area Under the Curve (AUC) of polysomes to that of 80S monosomes (**Figure 5E**), where SspH1 expression significantly reduced the polysome/80S ratio in wild-type and *gcn2* cells, but had no significant effect in *eap1* cells. These results confirm that Eap1, but not Gcn2, is required for the translational inhibition caused by SspH1, implicating the cap-dependent initiation machinery as a specific target of this effector.

## DISCUSSION

Bacterial pathogens and their hosts are locked in a continuous evolutionary arms race that drives reciprocal adaptations. While host organisms evolve intricate defense systems to detect and eliminate invading microbes, pathogens develop increasingly sophisticated strategies to counteract these defenses and ensure their survival. One of the most striking examples of such co-evolution is the ability of intracellular pathogens like *Salmonella enterica* to mimic eukaryotic enzymatic functions. Through the acquisition of E3 ubiquitin ligase activity, a hallmark of eukaryotic cellular regulation, *Salmonella* effectors have evolved to hijack one of the most central and versatile systems within the host, the ubiquitin-proteasome pathway, thereby manipulating cellular functions to their own benefit ^6,11^.

In this study, we provide compelling evidence that members of the NEL family E3 ubiquitin ligases from *Salmonella enterica* serovar Typhimurium, namely SlrP, SspH1, and SspH2, target a wide array of host proteins involved in fundamental cellular processes, with particular emphasis on those related to RNA metabolism, ribosome biogenesis and translation. Using ubiquitin enrichment proteomics in HEK293T cells, we identified over 200 potential substrates, with additional resolution obtained through the proteasome inhibitor MG132 (**Figure 1 and Table S1**). Our findings significantly expand the current repertoire of host targets and further support previous observations indicating that NEL effectors can manipulate not only immune signaling ^14,48^, and cell death regulation ^12^, but also other essential functions, such as translation, vesicle trafficking, cytoskeletal remodeling, and RNA splicing ^13^.

Gene ontology enrichment analysis revealed strong associations with RNA processing and translation, but also with actin cytoskeleton dynamics, post-translational protein modifications, chromatin remodeling, and regulation of cell death (**Figure 2**). These interactions suggest a broader, coordinated host subversion strategy employed by *Salmonella* through NEL ligases.

Interestingly, the observed enrichments align with previously reported individual targets and suggest a broader and coordinated strategy of host interference employed by *Salmonella* NEL ligases. Indeed, several targets and functional modules identified in this study overlap with datasets from published large-scale studies. For example, Walch *et al*. used proximity-labeling during *Salmonella* infection to uncover hundreds of host proteins interacting with effectors, including the NEL ligases ^24^. Notably, several proteins identified in our ubiquitylation dataset also appeared in their proximity network, suggesting physical and possibly functional engagement in the host. Additionally, while previous work by Fiskin et al. mapped global changes in the host ubiquitylome during *Salmonella* infection ^5^, revealing broad alterations across pathways such as RNA metabolism, cytoskeletal dynamics, and proteostasis, their study did not dissect the contribution of individual bacterial effectors. In contrast, our approach focuses specifically on the NEL family of E3 ligases and identifies putative substrates directly linked to their activity. Interestingly, several proteins identified in our screen (including UBE4B, DYNLL2, CLTA, and components of the spliceosome and ribosome), also appear in the ubiquitylation landscape described by Fiskin et al., suggesting that NEL effectors may play a significant and previously underappreciated role in shaping these infection-induced modifications ^5^.

Beyond global analyses, our proteomic dataset highlighted functional targets previously shown to be implicated in bacterial pathogenesis. Strikingly, several proteins involved in vesicular trafficking, including Rab14, R-Ras2, and phosphatidylinositol 4-kinase, were identified as putative ubiquitylation targets in our study. The manipulation of vesicular trafficking is a well-established strategy among *Salmonella* T3SS effectors. For instance, SopB modulates phosphoinositide signaling to promote membrane ruffling during bacterial entry ^49,50^, while SptP counteracts these changes to restore cytoskeletal activity post-invasion ^51^. SifA and SseG modulate the host endomembrane system and stabilize the *Salmonella* containing vacuole by interfering with Rab GTPases and microtubule-associated positioning ^52,53^. The identification of Rab14 and R-Ras2 in our screen suggests that NEL ligases may also contribute to this subversion of membrane trafficking, potentially via ubiquitin-mediated regulation of key trafficking hubs.

Two notable hits from our dataset (Larp4 and Larp7) were consistently identified across both experimental conditions. STRING-based network analysis (**Figure 2C, 2F**) placed them as central nodes within RNA processing and translation modules, respectively. Larp7 plays a key role in regulating transcription via the 7SK snRNP complex, which represses P-TEFb-dependent elongation. Interestingly, *Legionella pneumophila* targets Larp7 via its AnkH effector, disrupting host transcription ^30^, while *Coxiella burnetii* uses AnkG to similarly manipulate this complex ^54^. Our identification of Larp7 as a potential NEL effector substrate suggests that *Salmonella* may exploit the same node, through ubiquitylation, rather than via direct protein-protein interference. This convergence across phylogenetically distinct pathogens supports the idea that Larp7-dependent transcriptional regulation is a conserved vulnerability exploited during intracellular infection.

We independently validated a subset of candidates with high statistical confidence and novel biological implications, such as PRC1, DCD, NCLN, AURKAIP and SKP1 (**Figure 3**). This approach allowed us to explore effector impact beyond canonical immune targets, and to uncover new mechanistic avenues potentially unique to *Salmonella* pathogenesis.

Further insight emerged from analyzing protein-protein interactions involving our validated substrates. For example, ZC3H15, an interactor with NCLN, is upregulated during microsporidia infection ^55^, while POLDIP3, linked to our dataset, is a known target of *Ehrlichia chaffeensis* TRP47 effector ^56^. Moreover, eEF1A, which we identified in our dataset, is also targeted by *Legionella* Lgt1 to inhibit translation ^57^. It could be that eEF1A represents a common target for translation inhibition across intracellular pathogens. These overlaps not only validate our findings but also support broader hypotheses about convergent evolution in host manipulation. Our study also significantly extends these observations by uncovering dozens of new candidate substrates and suggests a multi-process subversion mechanism. Additionally, our strategy, using heterologous expression of effectors, allows finer resolution of effector-specific ubiquitylation events that are otherwise masked in whole-bacterium infection models.

One of the most notable outcomes of our study is the consistent enrichment of pathways associated with ribosome biogenesis and RNA processing (**Figure 1, 2A, 2B**). This is particularly timely in light of recent work emphasizing the role of ubiquitylation in regulating ribosome assembly and function ^25^. Functionally, we demonstrate that SspH1 impairs yeast growth (**Figure 4A**), as previously described by Keszei et al ^40^, and disrupts polysome profiles (**Figure 4F**), suggesting an impact in global translation as it is significantly impaired. Remarkably, the SspH1-dependent growth defect and translation inhibition observed in yeast is dependent on the eIF4E-associated protein Eap1 (**Figure 5A**), which is assumed to be a functional homolog of mammalian 4E-binding proteins ^46^. This places SspH1 within a broader group of bacterial effectors (e.g., *Shigella* IpaH and OspF^58^, *Legionella* SidI and Lgt1-3 ^57,59^) that modulate host protein synthesis. The mechanism by which Eap1 mediates the translation inhibition response exerted by SspH1 expression is currently unclear. Assuming the negative role of 4E-BPs in cap-dependent translation ^43^, we suggest that SspH1 ubiquitinates and likely degrades a factor required to maintain Eap1 activity repressed. Thus, the absence of Eap1 would restore global translation even in the presence of SspH1 (**Figure 5C**). Building on that evidence, we explored whether a related mechanism could operate in mammalian cells. To this end, we used the STRING database to identify human proteins from our NEL-ligase substrate dataset that potentially interact with eIF4E, a central regulator of cap-dependent translation. Notably, two RNA binding proteins, FXR1 and LSM14B, emerged as high confidence eIF4E interactors (**Figure S4**). FXR1 has been shown to facilitate eIF4E complex assembly and translation of AU-rich mRNAs by binding to 3’UTR elements ^60^, while LSM14B is involved in the regulation of mRNA storage and translation of developmental and stress context through ribonucleoprotein granule formation^61^. Importantly, FXR1 was previously identified as a SspH1 interactor in a proximity-labeling study conducted during *Salmonella* infection ^24^ providing independent support for a biologically relevant interaction. The identification of these proteins as putative SspH1 substrates raises the possibility that SspH1 may interfere with host translation by modulating the function or stability of FXR1 and/or LSM14B via ubiquitination. While further experiments will be required to confirm direct ubiquitination and clarify the downstream effects on host translational control, our data suggest that SspH1 could converge on the eIF4E/4E-BP regulatory axis through previously unrecognized nodes, adding another layer of complexity to its potential role in manipulating host gene expression.

A recent study by Wood et al ^62^ showed that *Salmonella* infection modulates host translation in macrophages through a mechanism independent of effector secretion, relying instead on the insertion of the SPI-1 T3SS apparatus. This process transiently enhances translation of transcriptional repressors like EGR1, dampening inflammation and promoting bacterial persistence. While these findings highlight an effector independent strategy, our study complement this by demonstrating that SspH1 modulates translation via the 4E-BP homologue Eap1 in yeast. Additionally, we identify other translation initiation factors (eg. EIF3 and EIF5) as well as a number of transcriptional regulators as potential substrates for the ubiquitin ligase activity of the *Salmonella* NEL effectors (**Table S1**). Overall, these findings suggest that *Salmonella* deploys both general and effector-specific mechanisms to reprogram host gene expression to their benefit.

Altogether, the multifunctionality of these NEL effectors underscores their role as powerful tools of host manipulation. The fact that SlrP, SspH1, and SspH2 converge on affecting translation, RNA biology, chromatin organization, and vesicular trafficking highlights the evolutionary pressure on *Salmonella* to control gene expression and protein turnover of its host. Importantly, these ligases differ in their specificity: while SspH1 and SspH2 are known to promote K48-linked degradation ^40,63^, SlrP may use non degradative chains such as K63 ^29^, allowing for more specific, non-canonical regulation. This functional divergence may reflect distinct evolutionary trajectories or cooperative strategies.

Our study is not without limitations. HEK293T cells, although ideal for transfection and proteomic analysis, do not fully recapitulate the complexity of natural infections. Moreover, simultaneous expression of all three effectors in a pooled format, while efficient, precludes assigning individual substrates with full confidence. Nevertheless, our approach enabled us to identify a diverse set of novel host targets, validate several interactions, and propose testable models for effector-specific functions.

Finally, we provide a public dataset of NEL-ubiquitylation substrates that may serve as a valuable resource in the field. Future efforts should focus on functional validation during infection, identification of specific ubiquitin linkages, and dissection of downstream consequences for signaling and immune defense of the host cell. Taken together, our findings reveal an unexpected level of complexity and versatility in *Salmonella* NEL ligases, expanding their role beyond immune modulation to include regulation of host gene expression, protein synthesis, and intracellular architecture. This work not only enhances our understanding of *Salmonella* pathogenesis but also contributes broadly to the study of bacterial effector-host interactions.

## ACKNOWLEDGMENTS

We would like to express our special thanks to Pedro Escoll for the scientific discussion that gave rise to the seminal idea for this work. We acknowledge Roberto Balbontín for the critical reading of the manuscript. We thank Ana Rodríguez Hortal, from the Mass Spectrometry laboratory at Pablo de Olavide University, for her support. This research is part of the project R+D+i PID2022-136863NB-I00 funded by MICIU/AEI/10.13039/501100011033 and by “ERDF a way of making Europe” to F.R.-M. and J. B.-B., and by the European Union’s Horizon 2020 research and innovation program under the Marie Skłodowska-Curie grant agreement No 842629 to J. B.-B. Work in J.d.l.C. laboratory is funded by the project R+D+i PID2022-136564NB-I00 from MICIU/AEI/10.13039/501100011033 and by “ERDF a way of making Europe”. J.d.l.C. also acknowledges the Andalusian Platform of Biomodels and Resources in Genomic Edition, the FORTALECE Program (FORT 2023) from MICU, and the Translacore (CA21154) and ProteoCure (CA20113) COST Actions from the EU (European Cooperation in Science and Technology) for support.

## AUTHOR CONTRIBUTION

Conceptualization: J.B.-B. and F.R.-M.; Investigation: A.B.-B, S.M.-V., C.V.G.; Data Curation: I.S.-F, F.P. and L.T.-G.; writing—original draft preparation, J.B.-B., I.S.-F., F.R.-M. and A.B.-B.; writing—review and editing, A.B.-B., I.S.-F., S.M.-V., C.G., F.P., J.d.l.C., L.T.- G., J.B.-B. and F.R.-M.; visualization, A.B.-B., J.B.-B. and I.S.-F.; funding acquisition, J.B.-B., F.R.-M. and J.d.l.C. All authors have read and agreed to the published version of the manuscript.

## MATERIALS AND METHODS

### Bacterial strains and growth conditions

*S.* Typhimurium ATCC 14028 was used as wild-type strain to amplify and clone the effectors into pcDNA3. Other bacterial strains used in this study are listed in Table S2. Bacterial cultures were grown at 37 °C with Lysogeny Broth medium (LB) supplemented when appropriated, with the following antibiotics: 50 μg mL^−1^ kanamycin (Kan), 100 μg mL^−1^ ampicillin (Amp). The solid media contained 1.5 % agar. For protein synthesis induction, isopropyl-β-D-thiogalactoside (IPTG) 1mM was added to the medium.

### Yeast strains and growth conditions

The *Saccharomyces cerevisiae* strains used are listed in **Table S2**. Yeasts were grown at 30 °C in YPD (1 % yeast extract, 2 % peptone, 2 % glucose) or synthetic drop-out (SD, 0.15 % yeast nitrogen base without amino acids and ammonium sulfate, 0.5 % ammonium sulfate, 2 % glucose, raffinose or galactose) medium. Yeast SD medium was prepared with Formedium supplements lacking leucine to select for the presence of derivatives of the pAS24 plasmid. The solid media contained 2 % agar. Yeast cells were transformed by the lithium acetate method ^64^. Yeast genetic techniques have been previously described ^65^.

### Mammalian cell culture

HEK293T (human embryonic kidney SV40 transformed; ECACC no. 12022001) cells were cultured in DMEM supplemented with 10 % fetal calf serum, 2 mM L-glutamine, 60 μg mL^-1^ penicillin and 100 μg mL^-1^ streptomycin. Cells were kept in a humidified atmosphere with 5 % CO_2_ at 37 °C. For transient transfection assays, Xfect reagent (Takara) was used according to the manufacturer’s instructions. For cell lysis, cells were incubated at 4 °C in NP40 buffer (10 mM Tris-HCl pH 7.4, 150 mM NaCl, 10 % glycerol, 1 % NP40, 1 % aprotinin, 1 mM PMSF, 1 μg mL^-1^ pepstatin and 1 μg mL^-1^ leupeptin) for 20 min. The extract was centrifuged at 12000 rpm for 20 min and the supernatant was stored at −80 °C.

### Plasmid construction

Plasmids are listed in **Table S2**. Amplification reactions were carried out on a T100 Thermal Cycler (Bio-Rad) using Q5 High-Fidelity DNA polymerase (New England Biolabs) or MyTaq Red DNA polymerase (Bioline) according to the supplier’s instructions. Oligonucleotides are described in **Table S2**. To generate pGEX derivative constructions and plasmids pIZ3717 and pIZ3737, vectors and inserts were PCR- amplified with appropriate overlapping edges and fused by Gibson assembly (Gibson et al, 2009). Overlapping PCR was used to generate point mutations in the gene encoding SspH1 using pIZ3717 as template. To generate pcDNA3 derivatives constructions and plasmids pIZ3728 and pIZ3731 the inserts were amplified by PCR and cloned into the vector by restriction enzyme digestion and ligation with the appropriate enzyme (New England Biolabs). Constructions were transformed into chemically competent DH5α cells and plated on LB-agar plates containing an appropriate antibiotic. Colony PCR was performed to check for the presence of the insert and identify positive clones. Single colonies were grown in 5 mL liquid LB medium at 37 °C overnight and the plasmid DNA was extracted using NucleoSpin® Plasmid preparation kit according to the manufacturer’s instructions (Macherey-Nagel). The plasmids were sequenced with an automated DNA sequencer (Stab Vida, Oeiras, Portugal).

### Enrichment of host-ubiquitylated proteins with UbiQapture-Q matrix

HEK293T cells were transiently transfected with SlrP, SspH1 and SspH2 expressing vectors (pIZ1725, pIZ3540, pIZ3541), or transfected with the empty vector. After 24 h, transfected cells were harvested, lysed and total protein was quantified using the Coomassie (Bradford) protein Assay Kit (Thermo Scientific) according to the manufacturer’s instructions. To avoid degradation of ubiquitylated proteins, some samples were treated with the proteasomal inhibitor MG132 (10 μM) during 6h before cells lysis. Upon quantification of total protein, the UBI-QAPTURE-Q kit (Enzo life Sciences) was used to perform a ubiquitylated proteins enrichment from each sample according to the manufacturer’s instructions with the following modifications. Total protein extracts were diluted 1:2 with PBS and mixed with 40 μL of matrix previously equilibrated with PBS. To favor binding of the proteins of interest to the matrix, samples were incubated at 4 °C overnight under agitation. The matrix was extensively washed with PBS and samples of the initial extract, the unbound fraction, the last wash and of the matrix-bound proteins were saved to check by western blot the presence of ubiquitylated proteins using the anti-ubiquitin HRP antibody (mono and polyubiquitinylated conjugates recombinant antibody (UBCJ2)) supplied with the kit. The matrix-bound proteins were boiled with Laemmli buffer with 100 μM DTT and stored at - 20 °C.

Three biological replicates of samples without MG132 treatment and 6 biological replicates of samples treated with MG132 were done.

### Mass-spectrometry identification of host ubiquitylated substrates

The samples were precipitated with methanol-chloroform, and the pellet was resuspended in 6 M urea (50 mM ammonium bicarbonate). Then,10 mM DTT and 30 mM iodoacetamide were added for reduction of disulfide bridges and carbamidomethylation of -SH residues, respectively. Samples were mixed with bovine trypsin (Promega) in a 1:12 ratio (enzyme: substrate) and incubated overnight at 37 °C. The reaction was stopped by adding formic acid at a final concentration of 0.5 %. Subsequently, C18-filled tips (OMIX, Agilent Technologies) were used to remove salts and concentrate the digested samples. The extracts were resuspended in 0.1 % formic acid for analysis by liquid chromatography-mass spectrometry.

Peptides were separated on an Easy n-LC 1200 chromatograph (Thermo Scientific) using a C18 Easy spray column (2 μm, 100 Å, 75 μm x 50 cm). The mobile phases used were water-0.1 % formic acid (phase A) and acetonitrile-20 % water-0.1 % formic acid (phase B). The separation was carried out maintaining a flow rate of 200 nL min^-1^ and applying the following phase gradient: 10 % - 35 % of phase B for 240 min, 35 % of B for 1 min and 100 % of B for 5 min.

The acquisition of spectra of the eluted peptides was carried out with an Orbitrap Q-Exactive Plus mass spectrometer (Thermo Scientific), fragmenting the 10 most intense peaks in each measurement cycle (DDA mode, top 10 MS/MS). The data obtained were analyzed with SEQUEST, a search engine of the Proteome Discoverer 2.2 software (Thermo Scientific). Cysteine carbamidomethylation was considered as fixed modifications, and methionine oxidation and lysine ubiquitylation were considered as variable modifications. The search was launched against the *Homo sapiens* database of Uniprot and a threshold FDR value of 1% was applied for the identifications obtained.

### Recombinant overexpression

*Escherichia coli* Rosetta containing pGEX-4T-1, pGEX-4T-2 derivatives and *E. coli* XL1-Blue, BL21 (DE3) or M15/pREP4 containing derivatives of pQE80L or pQE30, respectively, were grown in LB with appropriate antibiotic at 37 °C, and 200 rpm until the culture reached and OD_600_ of 0.5. Then 1 mM IPTG was added and the bacteria were grown at 30 °C, 200 rpm during 3 h. Bacteria were pelleted by centrifugation (8000 rpm, 10 min, 4 °C) and the pellets were resuspended in ice-cold NP40 buffer (for GST protein purifications) or imidazole lysis buffer (50 mM NaH_2_PO_4_, 300 mM NaCl, 10 mM imidazole) (for 6xHis protein purifications). The cells were subsequently disrupted using a sonicator (Branson 250) and the lysates were cleared by centrifugation step (12000 rpm, 20 min, 4 °C). Fusion proteins were isolated from bacterial lysates by affinity chromatography with glutathione-agarose beads (Sigma-Aldrich) or Ni-NTA agarose beads (Sigma-Aldrich), depending on the tag. 6His fusion proteins were eluted from the beads with imidazole elution buffer (50 mM NaH_2_PO_4_, 300 mM NaCl, 300 mM imidazole).

### Protein extraction, protein electrophoresis and immunoblotting

Bacterial and mammalian lysates were mixed with an equal volume of 4x Laemmli sample buffer and boiled for 5 min. Proteins were separated by SDS-PAGE using mini-protean TGX precast gels, 4-15 % gradient (BioRad). Proteins were transferred to nitrocellulose filters for western blot analysis using the Trans-Blot® Turbo™ Transfer System (BioRad) 30 min, 25 V, 1A. To block the membrane EveryBlot Blocking Buffer (BioRad) was used. Primary antibodies were anti-FLAG M2 (mouse, monoclonal, 1:5000, SigmaAldrich), anti-HA-peroxidase 3F10 (rat, 1:2000, Roche), anti-Mono and polyubiquitinylated conjugates recombinant (HRP conjugate, mouse, monoclonal, 1:1000, Enzo), anti-PGK1 (mouse, monoclonal, 1:10000, Invitrogen). Secondary antibody was goat anti-mouse IRDye 800CW-conjugated antibody (LI-COR). The bands were detected using the Odyssey Fc imaging system (LI-COR).

### In vitro ubiquitylation assays

Ubiquitylation reactions were performed in a 20 μL mixture containing buffer A (25 mM Tris-HCl, pH 7.5, 50 mM NaCl, 5 mM ATP, 10 mM MgCl_2_, 0.1 mM DTT), 0.25 μg of human recombinant E1 (Boston Biochem), 1 μg of E2 (human recombinant UbcH5b from Boston Biochem) and 1 μg of HA-tagged ubiquitin (Boston Biochem) in the presence or absence of purified 6xHis-effector and GST-substrate immobilized in a agarose-beads matrix. Reactions were incubated at 37 °C for 1 h at 1100 rpm in the Thermo Shaker Grant-bio PSC24N. The agarose-beads were extensively washed with NP40 buffer before the addition of an equal volume of Laemmli sample buffer. Specific substrate ubiquitylation was detected by immunoblot analysis using an anti-HA peroxidase antibody.

### Bioinformatics Data analysis of Proteomic Data

#### Data analysis and reproducibility

The quantitative data obtained from mass spectrometry was used to evaluate the reproducibility between biological replicates, Pearson’s correlation analysis was performed for each sample, as also for controls. Abundance non scaled values were transformed using Log_10_ prior to this analysis. Data analysis was conducted in Rstudio v2024.12.1 with Rbase v4.4.2. Individual correlation plots were generated using the corrplot package v0.95. Additionally, volcano plots visualizing protein enrichment (log_2_ fold change up or down 2.5 and p-value ≤ 0.05) were constructed with the ggplot2 package v3.5.1.

#### Protein network generation

Protein association networks were built using the Cytoscape software v3.10.3 (https://genome.cshlp.org/content/13/11/2498.full). For the network integrate interactions based on identified proteins the stringApp v2.2.0 (https://pubs.acs.org/doi/10.1021/acs.jproteome.2c00651) was used. Only the proteins found enriched in the sample were considered for the networks. Default parameters were generally applied, with the confidence cutoff set to 0.1 and the functional score cutoff set to 0.15. The proteins were colored according to the biological process and proteins of interest were highlighted manually.

#### Functional enrichment analysis

To identify overrepresented terms within the enriched identified proteins, we performed functional enrichment analysis. The ClueGO plugin v2.5.10 (https://pmc.ncbi.nlm.nih.gov/articles/PMC2666812/) within Cytoscape v3.10.3 environment was used. The default parameter was used with specific adjustments. GO-term fusion was selected, and only pathways with a p-value ≤ 0.5 after Benjamini-Hochberg correction were displayed. For analysis of biological processes and cellular components, the GO tree interval was set between 4 and 7, requiring at least 4 genes per term. For molecular function analysis, the GO tree interval was set between 1 and 10. For KEGG pathways, the Kappa score was set to 0.45 and at least 2 genes per term were required.

### Polysome Analysis and Sucrose Gradient Fractionation

Yeast extracts for polysome analysis were prepared and analyzed as previously described ^66,67^. Briefly, yeast was transformed with pAS24 derivatives constructs. Transformants were cultured in SD media with raffinose and without leucine until mid-log phase. Then, the cultures were diluted in SD with galactose and without leucine and harvested 16 h later. Cycloheximide was added to a final concentration of 100 μg mL^-1^ and incubated 15 min. Cells were pelleted by centrifugation at 5000 rpm in a microfuge during 5 min, at 4 °C, washed with polysome lysis buffer (10 mM Tris-HCl pH 7,5, 100 mM NaCl, 30 mM MgCl_2_, 100 μg mL^-1^ cycloheximide, 200 μg mL^-1^ heparin, 0,2 μL mL^-1^ DEPC) and centrifuged again. Cells were lysed by vigorously shaking with a vortex 8 min at 4 °C, in polysome lysis buffer with glass beads (425- 600 μm). Extracts were cleared by filtration and centrifugation at 15,000 rpm for 10 min at 4 °C. About 10 absorption units at 260 nm (A_260_) were loaded on the top layer of a 11 mL linear sucrose gradients (10 to 50 % prepared in 50 mM Tris-acetate (pH 7.5), 50 mM NH_4_Cl, 12 mM MgCl_2_, 1 mM DTT). These gradients were centrifuged at 39,000 rpm in a Beckman Coulter SW41Ti rotor for 2 h 45 min at 4 °C; the A_254_ was continuously monitored using an ISCO UA-6 system. GraphPad Prism software was used to quantify the polysome profiles by determining the area under the curve for each profile. The lowest absorbance value obtained for each profile was defined as the baseline. The peaks corresponding to each fraction were identified by equating the lowest point of each valley to the baseline value. The ratio of the area under the polysome curve to the area under the monosome curve (80S) was then compared between the different conditions.

### Data availability

The mass spectrometry proteomics data have been deposited to the ProteomeXchange Consortium via the PRIDE ^68^ partner repository with the dataset identifier PXD060966 and 10.6019/PXD060966.

## Supplementary Information

**Figure S1.**
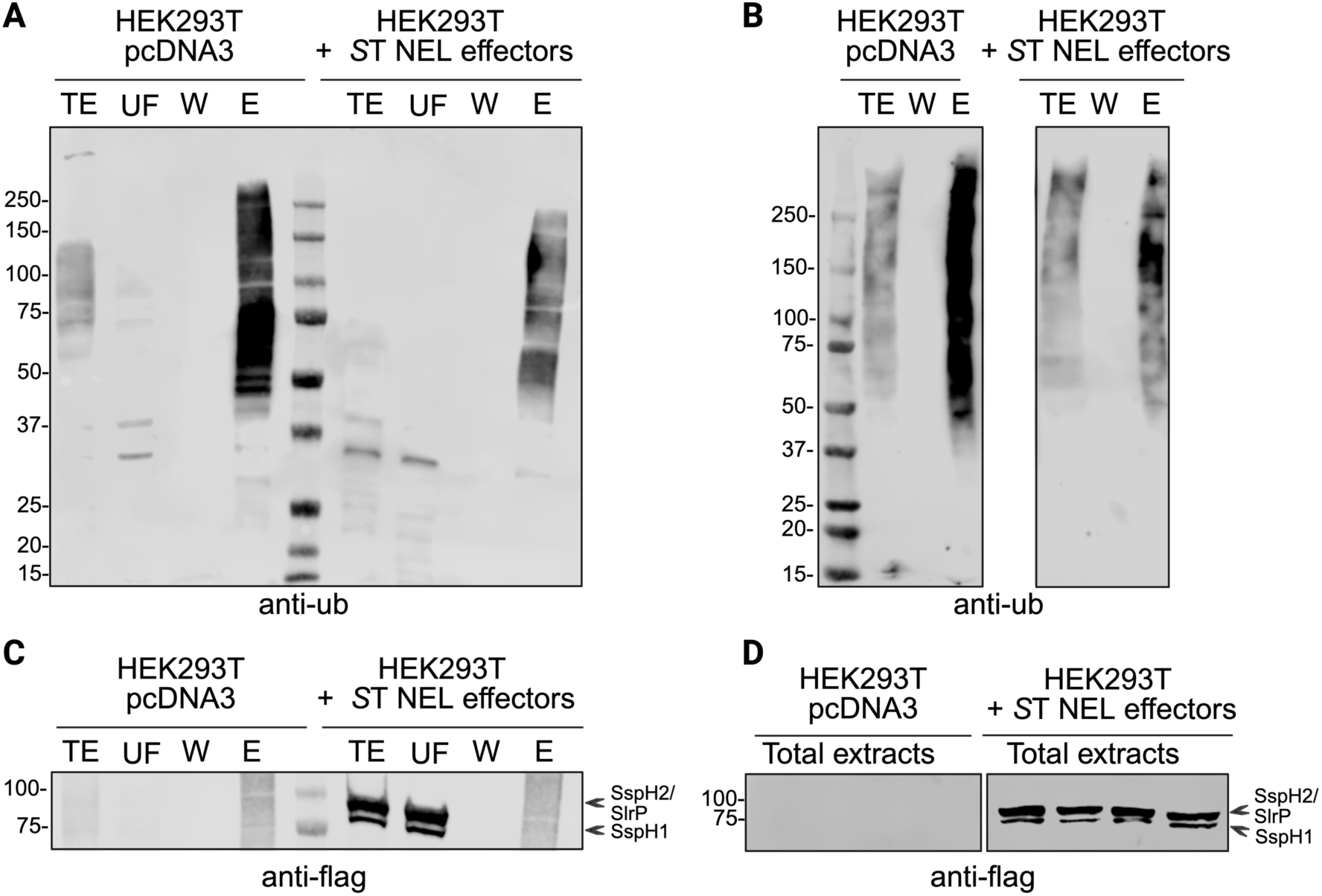
Enrichment of ubiquitin-conjugates proteins in HEK293T cells. Total cell extracts were subjected to mono- and poly-ubiquitylated proteins isolation using a high binding affinity matrix from the UBI-QAPTURE-Q kit (Enzo life Sciences). Samples of the total extract (TE), unbound fraction (UF), the last wash (W) and the proteins bound to the matrix (E) were loaded in SDS-PAGE and analyzed by immunoblot with anti-ubiquitin antibody (UBCJ2, Enzo). (A) Representative blot from n = 3 independent experiments performed with cells not treated with MG132. (B) Representative blot from n = 4 independent experiments performed with cells treated with MG132. To confirm the expression of the transfected bacterial effectors, immunoblots were performed using an anti-FLAG antibody in cells not treated (C) or treated (D) with MG132.

**Figure S2.**
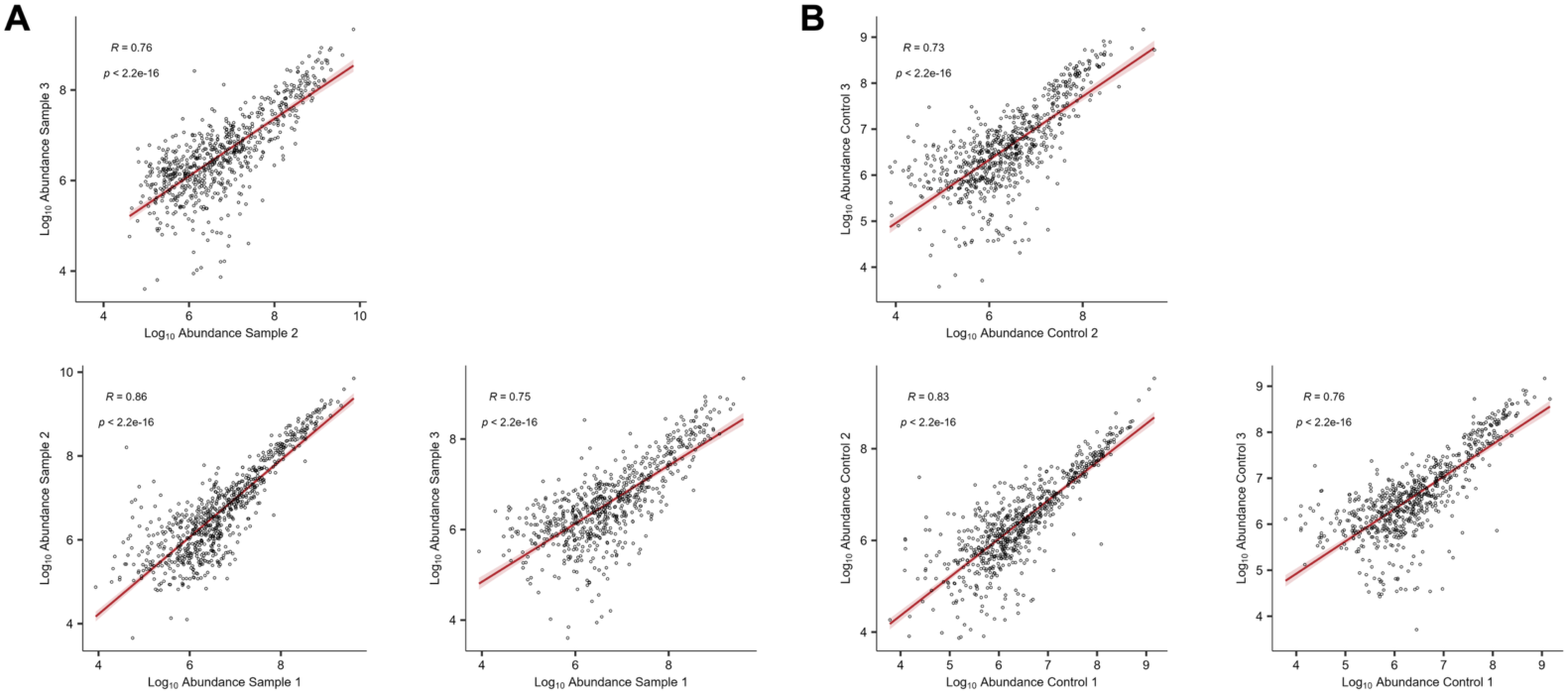
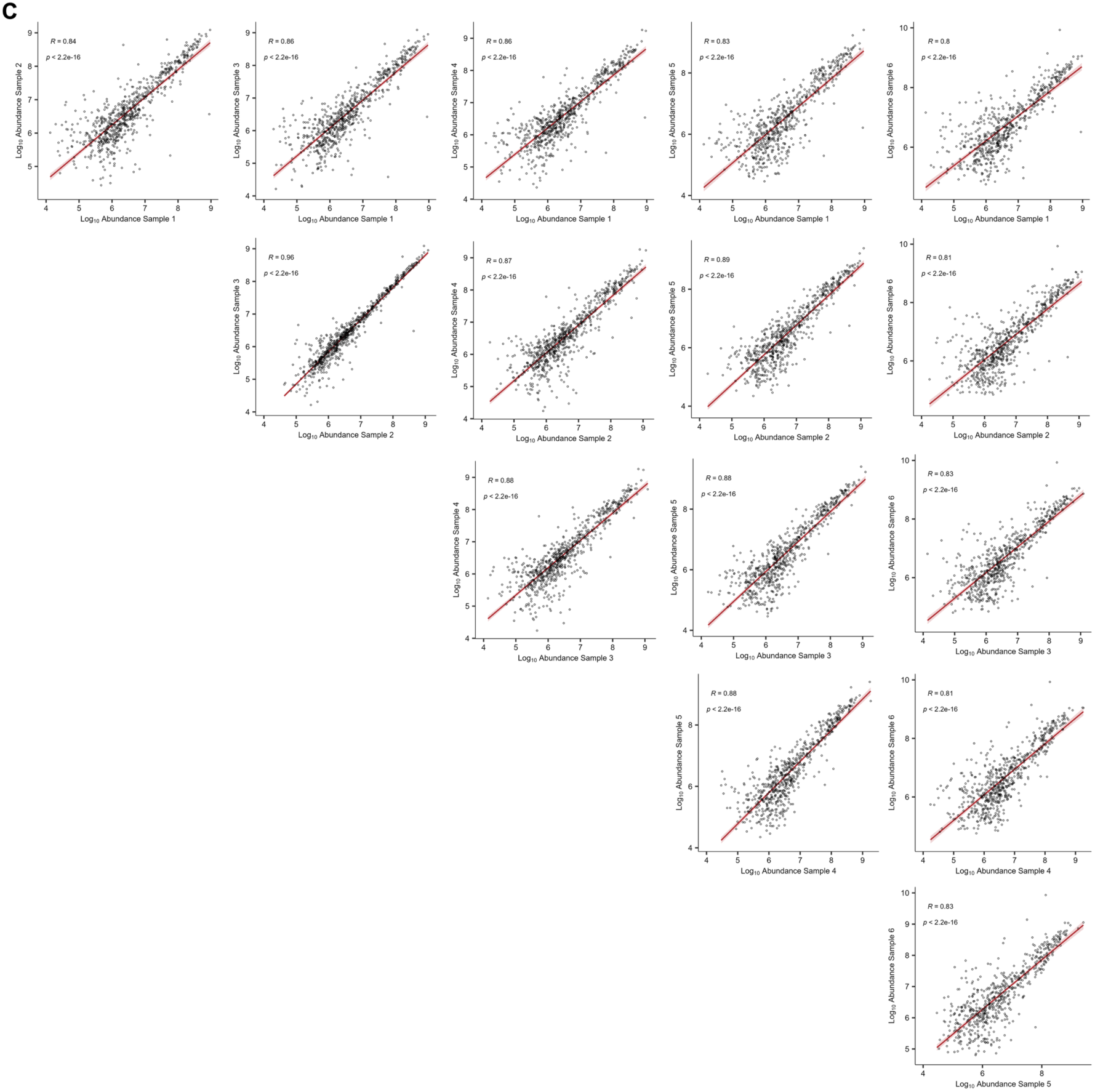
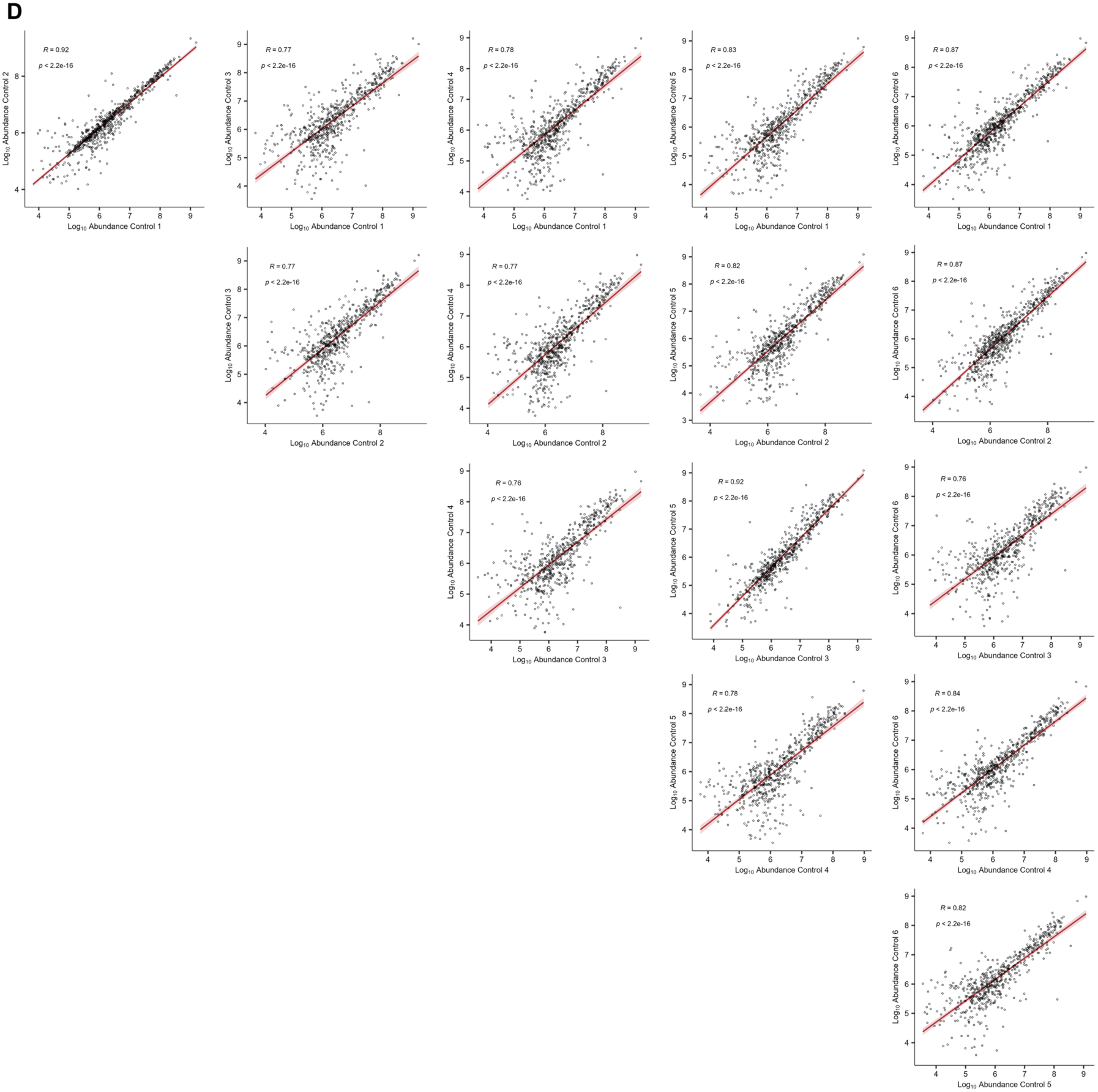
Reproducibility of proteomic data across biological replicates. Scatter plot of protein abundances of the samples -MG132 indicating Pearson correlation coefficients and the confidence intervals (A). Scatter plot of protein abundances of the controls -MG132 indicating Pearson correlation coefficients and the confidence intervals (B). Scatter plot of protein abundances of the samples +MG132 indicating Pearson correlation coefficients and the confidence intervals (C). Scatter plot of protein abundances of the controls +MG132 indicating Pearson correlation coefficients and the confidence intervals (D).

**Figure S3.**
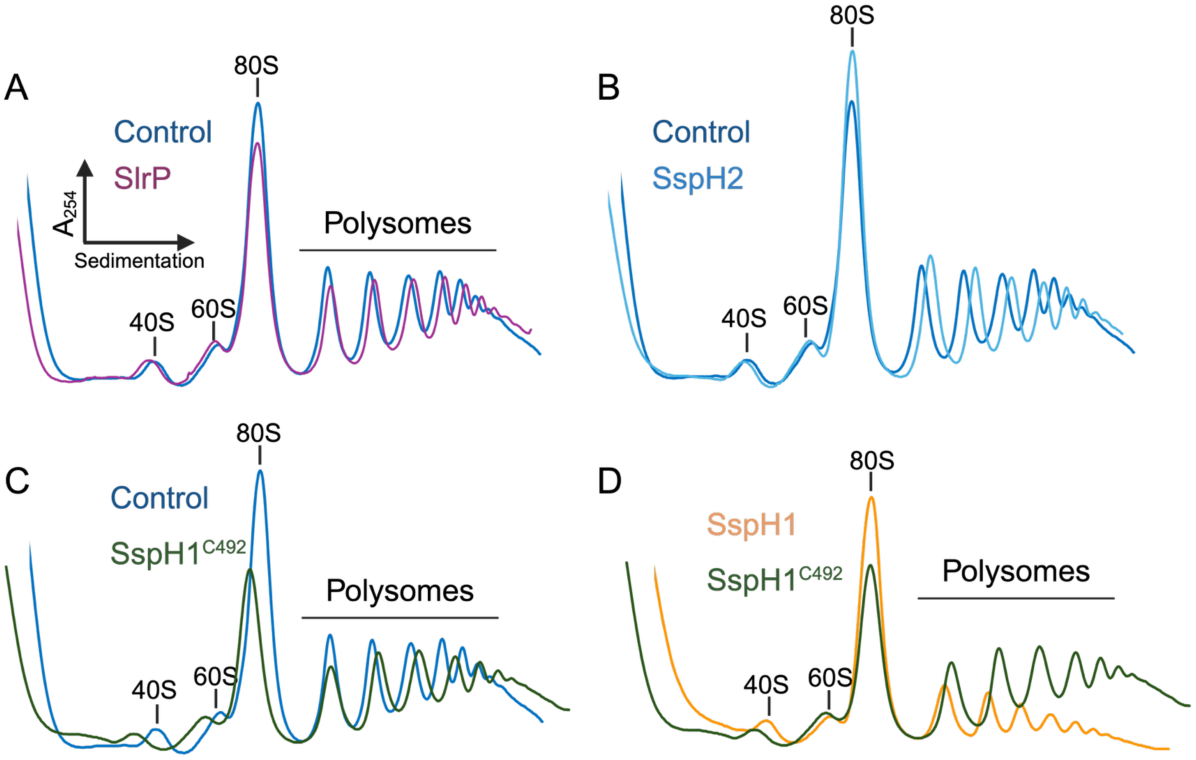
Polysome profiles of control cells and cells expressing SlrP, SspH2, SspH1, and SspH1^C492A^ effectors. Cells were lysed and polysomes profiles were performed as exactly described in M&M. Note that these profiles correspond to those shown in Figure 4. Translational differences between the samples were highlighted by aligning and merging the profile obtained for control cells with each of those obtained after expressing the different effectors SlrP (A), SspH2 (B), SspH1^C492A^ (C), as well as the comparison between SspH1 and the catalytic mutant (D).

**Figure S4.**
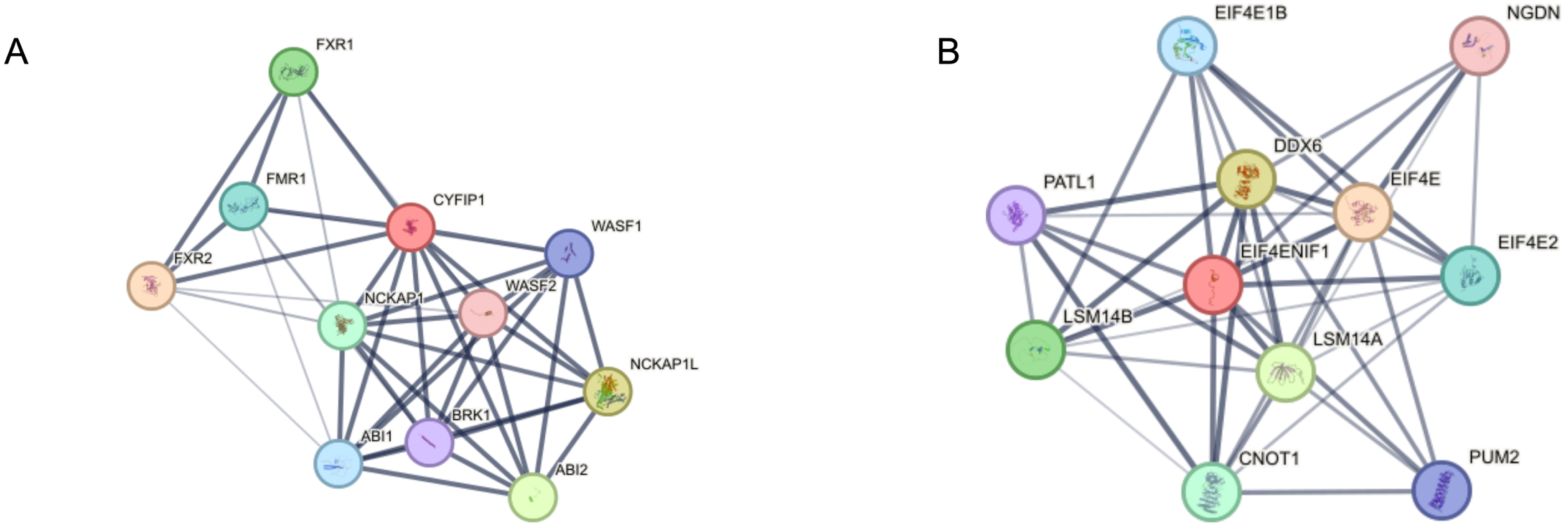
Protein-protein interaction network of FXR1 (A) and LSM14B (B) generated using STRING database. Network nodes represent proteins, with node color indicating the type of interactor: colored nodes represent query proteins and their first shell of interactors, filled nodes indicate proteins with known or predicted 3D structures. Edges represent protein-protein associations that are meant to be specific and meaningful (i.e., proteins jointly contribute to a shared function), though associations do not necessarily indicate direct physical binding. Edge thickness and color intensity indicate the strength of data support, with confidence scores ranging from low (0.150, thin lines) to highest (0.900, thick lines). Medium confidence is indicated by a score of 0.400, and high confidence by 0.700.

